# LAMA: Automated image analysis for developmental phenotyping of mouse embryos

**DOI:** 10.1101/2020.05.04.075853

**Authors:** Neil R. Horner, Shanmugasundaram Venkataraman, Ramón Casero, James M. Brown, Sara Johnson, Lydia Teboul, Sara Wells, Steve Brown, Henrik Westerberg, Ann-Marie Mallon

**Affiliations:** Medical Research Council Harwell Institute, Harwell, Oxfordshire OX11 0RD, UK; MRC Institute of Genetics and Molecular Medicine (IGMM), Edinburgh EH4 2XU; School of Computer Science, University of Lincoln, Lincoln LN6 7TS

## Abstract

Advanced 3D imaging modalities such as micro computed tomography (micro-CT), high resolution episcopic microscopy (HREM), and optical projection tomography (OPT) have been readily incorporated into high-throughput phenotyping pipelines, such as the International Mouse Phenotyping Consortium (IMPC). Such modalities generate large volumes of raw data that cannot be immediately harnessed without significant resources of manpower and expertise. Thus, rapid automated analysis and annotation is critical to ensure that 3D imaging data is able to be integrated with other multi-dimensional phenotyping data. To this end, we present an automated computational mouse phenotyping pipeline called LAMA, based on image registration, which requires minimal technical expertise and human input to use. Designed predominantly for developmental biologists, our software performs image pre-processing, registration, statistical and gene function annotation, and segmentation of 3D micro-CT data. We address several limitations of current methods and create an easy to use, fast solution application for mouse embryo phenotyping. We also present a highly granular, novel anatomical E14.5 (14.5 days post coitus) atlas of a population average that integrates with our pipeline to allow a range of dysmorphologies to be automatically annotated as well as results from the validation of the pipeline.

## Introduction

A major goal in biomedical research is to assign functional roles to all genes in order to shed light on disease mechanisms, and to identify disease-associated genes and novel drug targets. However, almost two decades since the human and mouse draft genomes were published (Lander et al., 2001; Waterston et al., 2002), the proportion of genes still in the dark genome (minimal gene function or disease association assigned) remains at over 30% (Oprea 2019). One important approach to add functional knowledge to each gene in the genome involves the phenotyping of mutant mice, leveraging the close homology between the mouse and human genomes as well as the relative ease of generating null deletions and the existence of established phenotyping procedures (Hrabě de Angelis 2015; Meehan et al., 2017). The International Mouse Phenotyping Consortium (IMPC) is a high-throughput functional genomics project that is well on its way to generating a genome wide catalogue of gene function by phenotyping all gene knockouts in mice on a uniform genetic background (Brown et al., 2012; Lloyd et al., 2020). To date phenotype data for over 7000 genes are available on the IMPC web portal (mousephenotype.org), data which has already contributed to the identification of many novel candidate disease genes and potential new mouse models of human disease (Cacheiro et al., 2020; Meehan 2017; Bowl et al., 2017; Moore et al., 2018).

Approximately one third of all knockout mouse lines exhibit embryonic or early postnatal, lethality or subviability (Hrabě de Angelis et al., 2015; Dickinson et al., 2016). These genes are of particular importance as they can provide insights into developmental processes and disorders. The IMPC embryo phenotyping pipeline aims to study these genes at key developmental time points, with a major component being the generation and analysis of high resolution, whole embryo, 3D images (Adams et al., 2013).

Embryo anatomy can be manually assigned phenotype annotation by experts with knowledge of mouse anatomy and development (Wilson et al. 2016). However, this is prohibitively time consuming to perform at scale and is subject to user error and bias. Furthermore, manual annotation is susceptible to inter-operator variability, and may miss abnormalities that are not present on a predetermined checklist of structures that reflect annotator expertise. Automated analysis methods that aim to address these issues have been developed and were first applied to mouse whole embryos, by analysing magnetic resonance images (MRI) from wild type mice at embryonic day 15.5 (E15.5) (Zamyadi et al. 2010). This work was expanded to the analysis of micro-CT images at E15.5 and incorporated the analysis of mutant embryo images (Wong et al., 2014). The latter works by aligning with a non-affine spatial transformation the gross anatomy of mutant embryos that share a genetic alteration and wild type embryos, and then localizing regions of anatomical dysmorphology by modelling the effect of genotype on local volume changes or voxel intensities, in a process called voxel-based morphometry (VBM). The resulting heat maps, representing statistical significance of local voxel deformation, can be overlaid on embryo images to highlight local dysmorphology. Mouse embryo atlases in which visible anatomical structures have been delineated enable the assignment of dysmorphology to anatomical labels as well as the automatic calculation of organ volume by the propagation of atlas labels post-registration back towards the input images. Atlases can be made using a single segmented specimen as a reference, for example those made for the eMouse Atlas Project (EMAP), which created atlases from multiple developmental stages from serial histological sections (Richardson et al., 2014). However the segmentation of organs from micro-CT images and MRI can be more difficuklt due to increased noise and reduced contrast. A solution to this is to segment a population average consensus image formed from spatially normalising multiple specimens, as shown by (Holmes et al., 1998) in brain MRI, and provides an unbiased registration target from which to propagate anatomical labels (Hammers et al., 2002). Mouse embryo atlases of population average models derived from MRI images (Cleary et al., 2010) and micro-CT (Wong et al., 2012) have been developed for the E15.5 stage where 6 and 48 structures were segmented respectively. These methods have yet to be applied to the analysis of other key developmental stages that are studied within the IMPC, namely E14.5, at which point organogenesis becomes largely compete, and E18.5, the stage immediately prior to birth and which is an important stage for the analysis of perinatal lethality or subviability.

Sample size of both mutant and wild type controls will have a substantial effect on the power of a mouse phenotyping study and its consideration in automated analysis of mouse embryos has not been investigated in-depth. Wong et al., (2012) performed a power analysis using automatically-calculated wild type organ size distributions from four organs (brain, liver, lung and myocardium), and showed that organ volume differences between 9-14% were theoretically detectable using a sample size of eight mutants and controls. Their analysis also showed that increasing sample size may substantially improve sensitivity, but they chose a sample size that was a compromise between sensitivity and computational expense imposed by the analysis pipeline design. Up to now, however, no empirical assessment of the effect of sample size has been carried out on whole embryo data. Imaging data submitted to the IMPC embryo pipeline often have mutant sample sizes that are significantly lower that the desired eight mutant specimens due to cost and other practical limitations. However, imaging data from wild type control specimens can be much higher as they accumulate over the lifetime of a project. Applying these large numbers of wild type control animals in the automated analysis should increase statistical power, but this has yet to be investigated.

A surprising finding from recent high-throughput studies is that mutant lines generated from isogenic inbred mice frequently exhibit incomplete penetrance and variable expressivity of phenotypes (Wilson et al., 2017; Dickinson et al., 2016). These phenomena potentially complicate efforts to assign phenotype annotations at the line level. The ability to reliably assign phenotype annotations to each animal (specimen-level annotation) would help with cases of penetrance and expressivity variability. In this work we address this question by empirically testing the ability to detect sex differences and phenotypes of knockouts with varying sample sizes.

Another factor to consider is the developmental stage of specimens being tested, as it is known that embryos harvested at E14.5 represent a range of developmental substages due to uncertainty around the exact time of conception. It has been shown that selecting wild type comparison specimens that are stage-matched is vital to prevent misidentification of dysmorphology during manual annotation of embryos (Geyer *et al.*, 2017), but this problem has yet to be studied in the context of automated phenotyping using 3D images.

The issues raised above motivated us to develop a new automated phenotyping pipeline, which we present in this paper. The software pipeline ‘Lightweight Analysis of Morphological Abnormalities’ (LAMA) is freely available software developed for Linux and Windows™ platforms (https://github.com/mpi2/LAMA). LAMA is an automated computational phenotyping pipeline based on image registration that performs image pre-processing, registration, statistical annotation and segmentation of 3D image data. LAMA requires minimal technical expertise to install, and can run on a standard desktop PC. The registration strategy that we have employed speeds up the spatial normalization process and enables a greatly increased sample number of baselines to be used as controls, thereby increasing the power to detect abnormalities. We also present results of the validation of the pipeline by testing the ability of LAMA to detect sex-specific differences between sets of E14.5 embryos as well as showing that it can uncover known anatomical abnormalities in a broadly affected knockout line as well as a knockout line displaying specific, localised dysmorphology. This represents the first time that VBM methods have been applied successfully to whole embryos at this developmental stage. We also show that the developmental substage is an important variable when designing automated phenotyping experiments and we provide a solution to mitigate this confounding effect. We have begun to address the issue of variable expressivity and incomplete penetrance and discuss future areas of research in these domains.

To accompany the pipeline, a novel, highly detailed E14.5 anatomical atlas of a population average is also presented. The population average was computed from micro-CT images of 16 wild type mice at a high resolution of 14 µm^3^. The atlas consists of the manual segmentation of 184 organs and anatomical structures in the population average representing the most detailed average mouse embryo atlas currently available. The population average and atlas will be a valuable resource for the developmental biology community to aid in both manual and automated analyses. Using them along with LAMA, it is possible for the first time to automatically annotate dysmorphic E14.5 embryo anatomy with ontological EMAPA terms (Hayamizu et al. 2013), which are ready for export to the IMPC for display on the portal, and as we move forward, this novel dataset of ontological phenotype associations will be integrated with other data sets enabling to study the gene function relationships to anatomy and ultimately disease.

## Results

### Overview of the phenotyping pipeline for E14.5 mouse embryos

LAMA is a voxel-based morphometry approach to automate the detection of anatomical dysmorphology in mouse embryos. It is written in the Python programming language and features spatial normalisation of images using a groupwise registration process to iteratively align micro-CT embryo images into the same coordinate space. Internally, it uses elastix (Klein et al., 2010) for pairwise multi-resolution 3D image registration. Firstly, a population average model embryo must be constructed from wild type derived images (Fig 1A), which serves as an atlas template due to its high contrast and low signal to noise ratio. The population average also acts as a target image for the subsequent spatial normalisation of the phenotyping images (described below) (Fig 1A). The next step involves spatially normalising each wild type and mutant image that will be used in the downstream statistical analysis by registering it onto the population average (Fig 1B), resulting in homologous anatomical structures occupying identical coordinates and morphology. After spatial normalisation, applying the inverse registration transformation to the population average mask and atlas, aligns them back onto the corresponding specimen image and allows for the calculation of whole embryo volume and each of the atlas labels (Fig 1B). For this paper, we focus on the organ volume analysis as this is linked to the atlas resulting in automated EMAPA-associated phenotype calls that are more robust and interpretable than those at the individual voxel level. However, we also include statistical parametric heat maps from the voxel-level Jacobian determinants (which indicate local volume shrinkage/expansion during spatial normalisation) for illustrative purposes.

**Figure 1.**
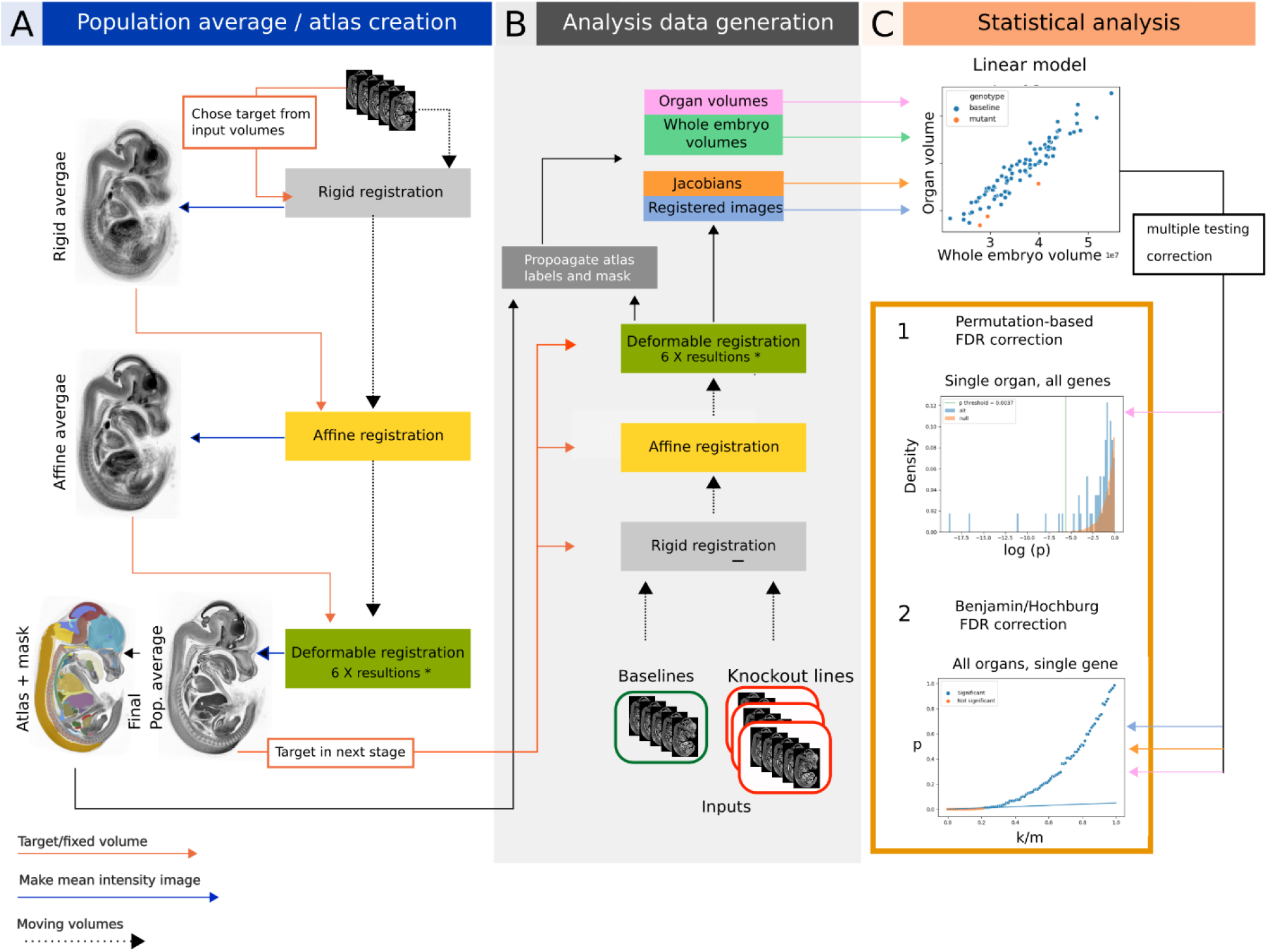
LAMA pipeline workflow. (A) Population average construction. An initial target from the inputs is used to rigidly align all inputs creating a rigid population average. This is repeated with affine registration and deformable registration at each stage using the population average as the target from the subsequent stage. (B) Generation of data for phenotype detection. The same process as in A, but each stage uses the population average form A as the target. Jacobian determinant volumes, registered images and organ volumes are statistically analysed. (C) Organ volumes (shown here) or voxel values are fitted to a linear model by genotype and whole embryo volume. The resulting genotype effect p-values are corrected for multiple testing either by permutation-based FDR correction (option 1, organ volumes only) the orange histogram is the null distribution from permuting the wild type organ volumes and the blue histogram is the alternative distribution derived from testing the organ volume from all mutant lines tested. The vertical green line indicates the calculated p-value threshold for this organ, with values lower than this annotated as significant. Option 2 is to apply Benjamini Hochberg FDR correction for voxel-level or organ volume data. k/m = p-value rank divided by number of values. The straight blue line indicates the threshold under which values are annotated as significant.

### Creation of a Novel E14.5 mouse embryo atlas

The E14.5 population average used in this study was created from 16 specimens (8 male and 8 female), with a resulting crown-rump length of 9.18 mm and an isotropic image resolution of 14 μm^3^ (Fig. 2A). The visible organs were segmented using a mixture of manual and semi-automatic segmentation (see Methods) producing a segmented E14.5 atlas containing 184 unique labels (Fig. 2B; Table 1). A key requirement for the full utilization of the atlas, is that the segmented labels are assigned appropriate community-accepted ontology terms allowing the automatic integration of the resulting gene-to-phenotype data from LAMA with other data sets. The 184 labels were therefore mapped to gross anatomy terms from the EMAPA developmental ontology (Hyamizu at. al., 2013). Through manual inspection of registration results a number of the labels were identified as being too small or thin leading to difficulty in assessing registration accuracy, such as the external carotid arteries (Fig. 2C). With this in mind, we identified such labels in the atlas (see Methods) to exclude them from downstream analyses, resulting in a final set of 103 labels that were used in the current analysis (Table 1). These 103 labels were distributed across the majority of the EMAPA high-level organ system terms (Fig. 2E) and range in size from the largest (forebrain at 35,596,092 μm^3^) to smallest (metatarsals 14,420 μm^3^) (Fig. 2D).

**Table 1.**
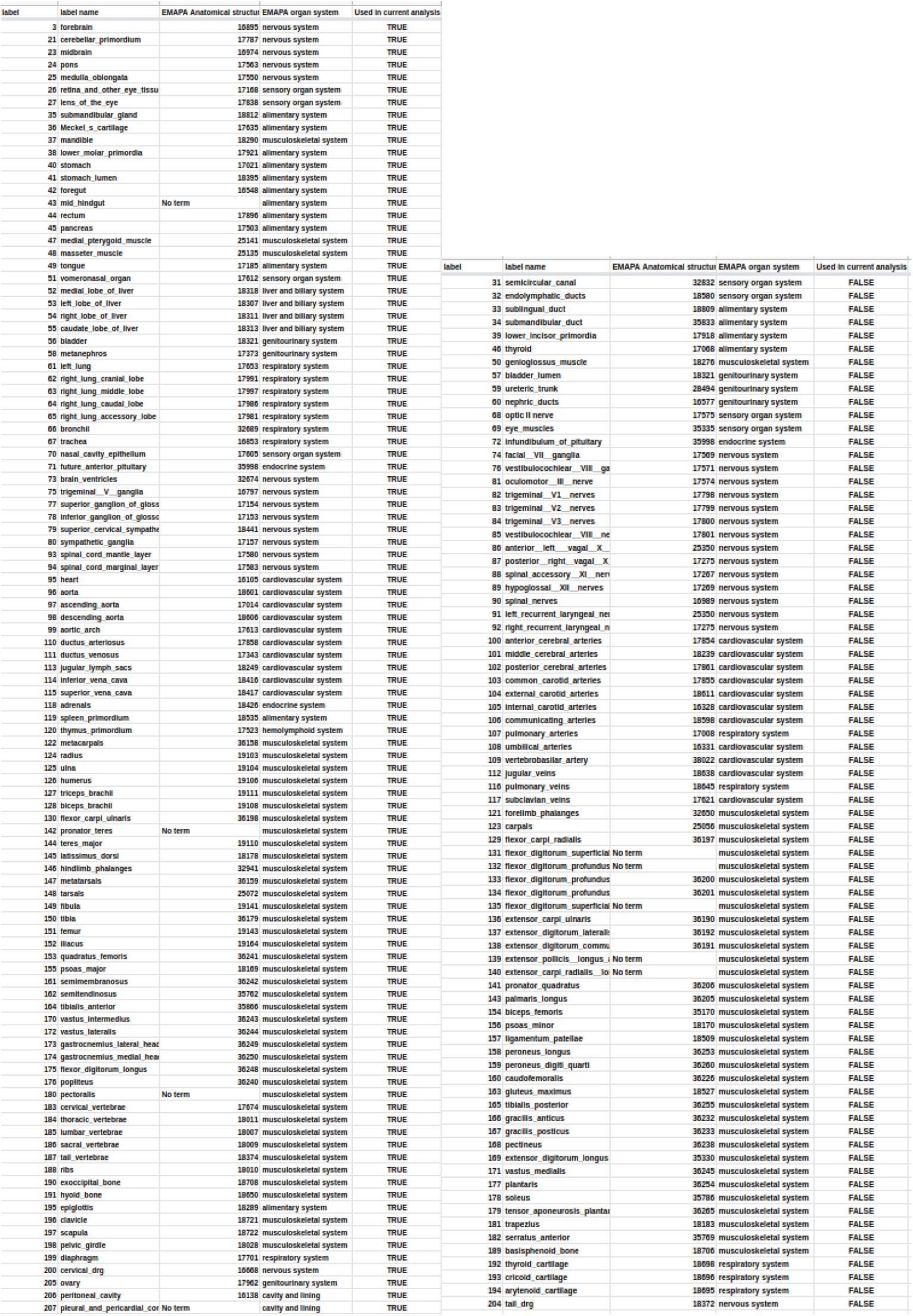
E14.5 Atlas label information label: label number in atlas images, label_name: the descriptive name, EMAPA Anatomical structure: associated EMAPA anatomical term, EMAPA organ system: the top level EMAP organ system terms, Used in current analysis: If FALSE not used in the current study.

**Figure 2.**
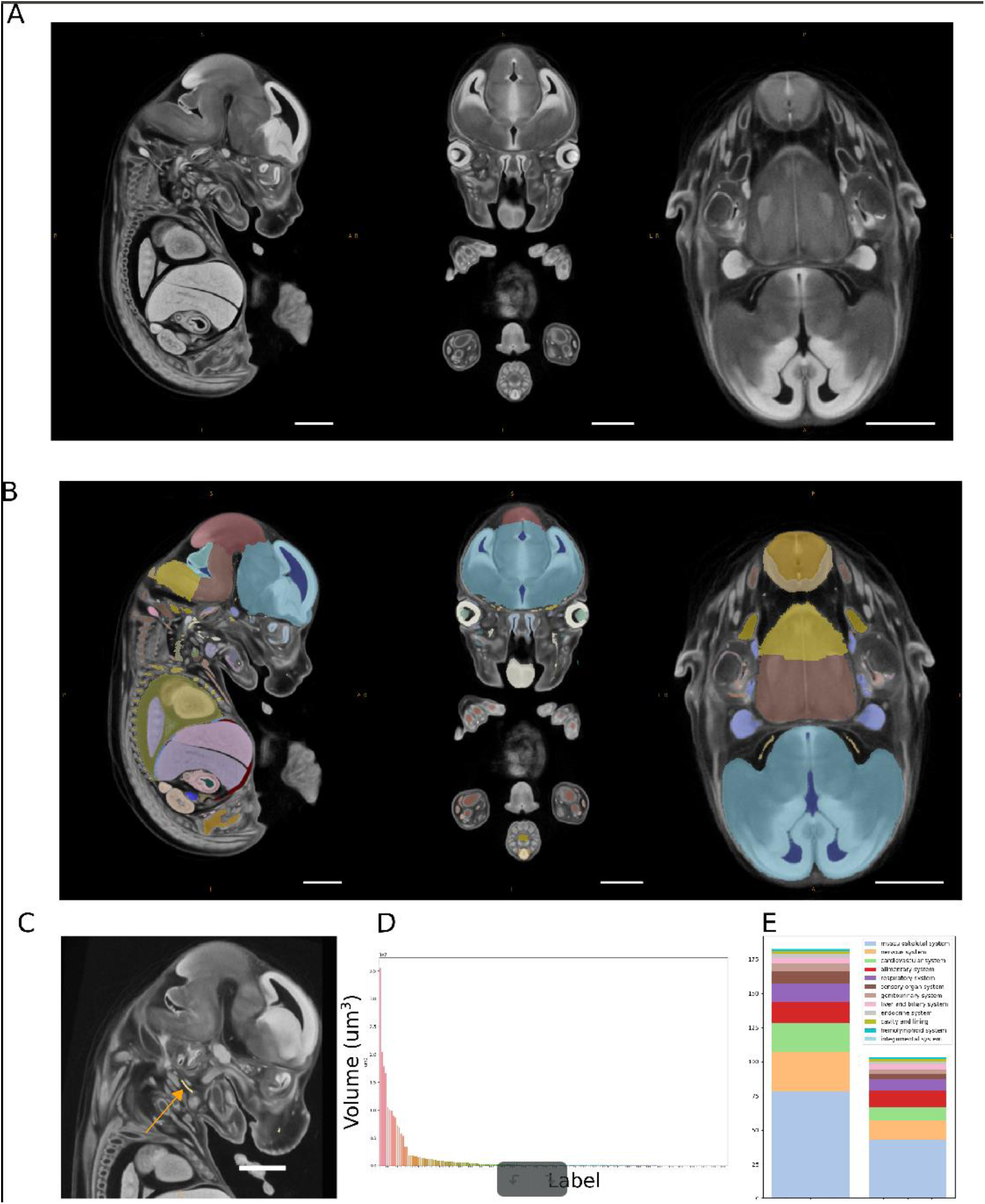
E14.5 atlas creation. (A) Population average created from 16 wild type mixed sex E14.5 C57BL/6 mice. (B) E14.5 atlas consisting of 184 individual structures overlaid on the population average. (C) An example of a small size-excluded label, external carotid artery (arrow). Scale bar = 1mm. (D) Plot showing the organ volumes for each of the 184 E14.5 atlas labels. (E) Counts for each high level organ system applied to labels in the E14.5 atlas.

### Developmental substage

Embryos harvested at E14.5 represent a range of developmental substages (DSS) and have rapidly developing anatomy and so it is crucial for a high throughput data analysis pipeline to account for this variation in the data. To account for overall size of the embryo, derived organ volumes are normalised by whole embryo volume and the Jacobian determinants are normalised by limiting the analysis of the Jacobian determinants to those generated after the affine registration stage, which accounts for overall embryo size (see Methods) as done previously at E15.5 (Wong et al., 2012). To assess whether this normalization is sufficient to account for differences due to embryo DSS, we made two datasets comprising of wild type specimens where eight of the smallest or largest specimens were labelled as “mutants” (see Methods), thereby simulating the cases of mutant lines containing embryos at early or late E14.5 DSS and we applied the LAMA pipeline statistical analysis to each data set. (Fig 3A). Both tests returned significant Jacobian determinant voxels suggesting that the relabelled wild types had morphological differences that were dependent of the developmental stage of the specimens. We then included a surrogate for DSS, whole embryo volume (WEV), within our linear model analysis, which resulted in the two tests returning no significant voxels for genotype effect, showing that the DSS effect can be controlled for in this manner. To gain a more detailed view on the voxel-level DSS-dependent relative size differences, we fitted Jacobian determinates from 93 wild type specimens to a linear model by WEV only (Fig 3B) which highlighted regions that are proportionally larger at later stages (red) or proportionally smaller at later stages (blue). Similarly, the stage effect on WEV-normalised organ volumes was significant for 78/103 labels (Table 2) including organs that were proportionally larger (n=58) later in development such as thymus and lung lobes (Fig. 3C-D) and those that were proportionally smaller (n=23) including brain ventricles and trigeminal glands (Fig. 3E-F). To summarize, normalizing the Jacobian determinants or organ volumes before statistical analysis is not sufficient to account for DSS and failure to model DSS can lead to false positive results, and one way to account for this is to regress out the DSS effect in the statistical analysis.

**Table 2.**
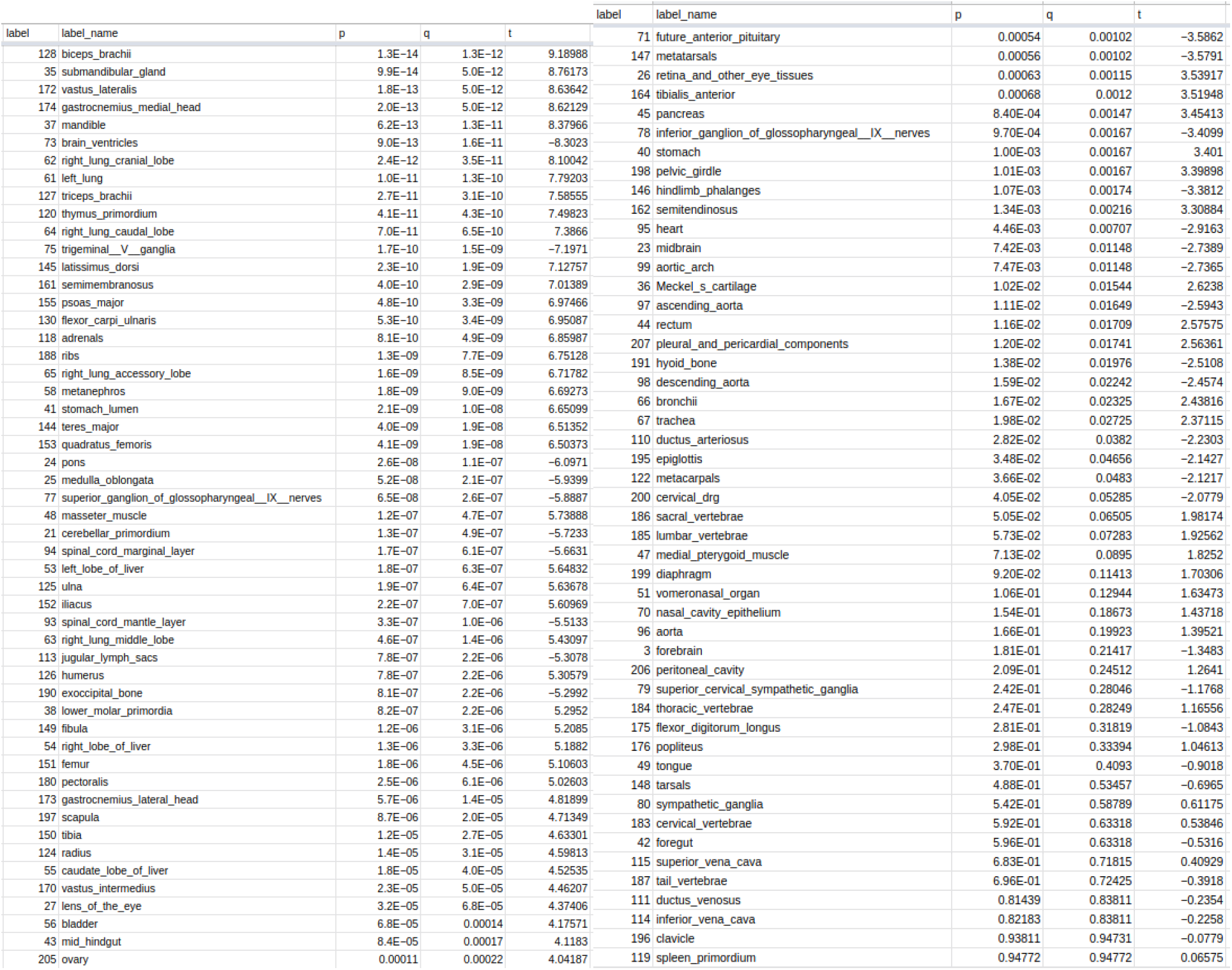
Wild type staging effect linear model outputs. Results from fitting calculated organ volumes to linear model : organ volume ∼ whole embryo volume (WEV). p: WEV effect p-value. q: Benjamini-Hochburg FDR-corrected p-values, t: WEV effect t-statistics.

**Figure 3.**
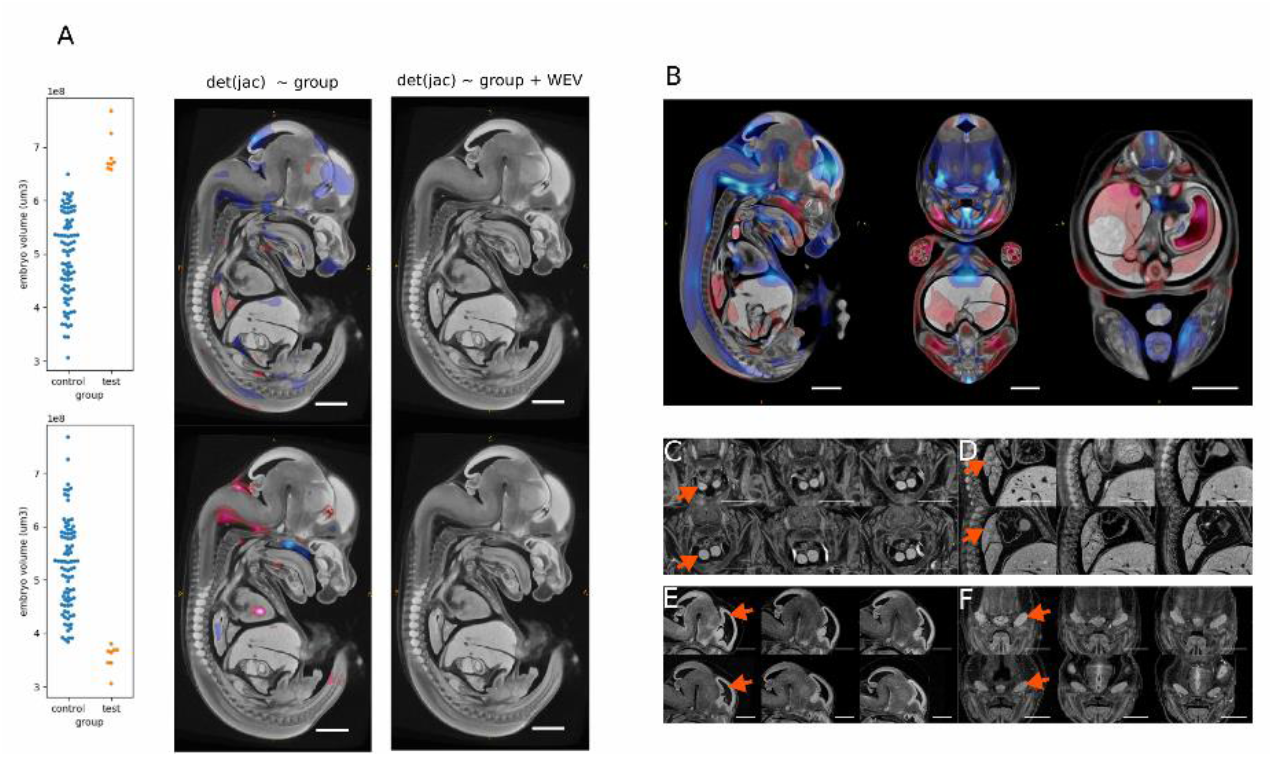
Effect of E14.5 developmental substage on local volume changes detected by LAMA. (A) Simulation of the analysis of mutant lines with large (top panel) or small (bottom panel) wild types relabelled as mutants. Genotype effect t-statistics where q (FDR-corrected p-values) < 0.05 were overlaid on the E14.5 population average. Left panel shows results from the model: *Jacobian determinant ∼ genotype* with red voxels highlighting regions that are significantly larger in the test group and blue voxels highlighting regions that are significantly smaller in the test group. Right panel shows results from model: *Jacobian determinant ∼ genotype + WEV*, where no voxels passed the q < 0.05 threshold. (B) Result from linear model: *Jacobian determinant ∼ WEV*, indicating regions where the normalised organ volume is directly (red) or inversely proportional (blue) to WEV. (C-F) Illustrative examples of differences in organ size relative to embryo volume. Each panel shows three representative wild types from the smallest set (top) and largest set (bottom) of specimens. Images are affinely registered towards the population average to account for overall embryo volume. Arrows indicate relevant anatomy. (C-D) Larger relative thymus size and lung volume (D), (E-F) Smaller relative lateral ventricles and trigeminal gland volume. Scale bar = 1mm.

### Optimal sample size for phenodeviance testing

In order to validate LAMA with a positive control we applied it to wild type embryos where females were relabelled as “mutant”. This gives us a convenient dataset where we expected the only gross differences between the sets to be located at the gonads (see Fig. 4, suppl. 1 for an example of gonad images). A quality control step removed specimens with obvious tissue damage due to sample preparation, extreme imaging artefacts or non-determined sex, resulting in a data set consisting of 89 wild type specimens (49 male, 40 female). Using a linear model (organ volume ∼ sex + whole embryo volume), we found that along with the gonad (FDR-corrected q-value = 1.3e^-25^) (Fig. 4A), the lens of the eye unexpectedly also had a significant sex effect (FDR-corrected q-value = 0.028) (Fig. 4B; Suppl. Fig 4,2). Therefore, we included the lens along with the gonad as true positives in the following experiment. With this large data set, we were also able to address the effect that sample size has on the sensitivity of phenodeviance detection (in this case the ability to differentiate between male and female gonads and lenses). We ran a series of experiments in which organ volume was first normalised by whole embryo volume and then regressed on whole embryo volume and sex as described previously, in which each experiment had a different number of male and female (relabelled as “mutant”) specimens. Each experiment was repeated 50 times with random specimen selection, and using the permutation approach to correct for multiple testing (see Methods). Due to the large difference between male and female gonad sizes, significant gonad volume differences were identified in almost every replication of each experiment (Fig. 4C), and significant Jacobian determinant voxels were identified within, or close to, the gonad with significant voxels covering a larger area with increasing male sample size (Fig 4E). For the lens of the eye, significant volume differences were detected only with a male sample size of 32 or over, with the maximum male and female sample size (49 male & 8 female) resulting in significant hits in over a half the cases (Fig. 4D). To assess the rate of false positive detection, any significant organ volumes, other than gonad or lens of the eye in the preceding experiments were classed as false positives. The rate of false positives was found to be well controlled with only 1 of 103 organs called as significant in more than 1% of tests (epiglottis in 1.6% of all the replications), and with a mean false positve rate of 0.07% per label. These experiments show that LAMA is able to identify sex-specific differences in wild type embryos and that even with a low mutant sample size, differences in morphology can be detected more reliably as the control sample size is increased.

**Figure 4.**
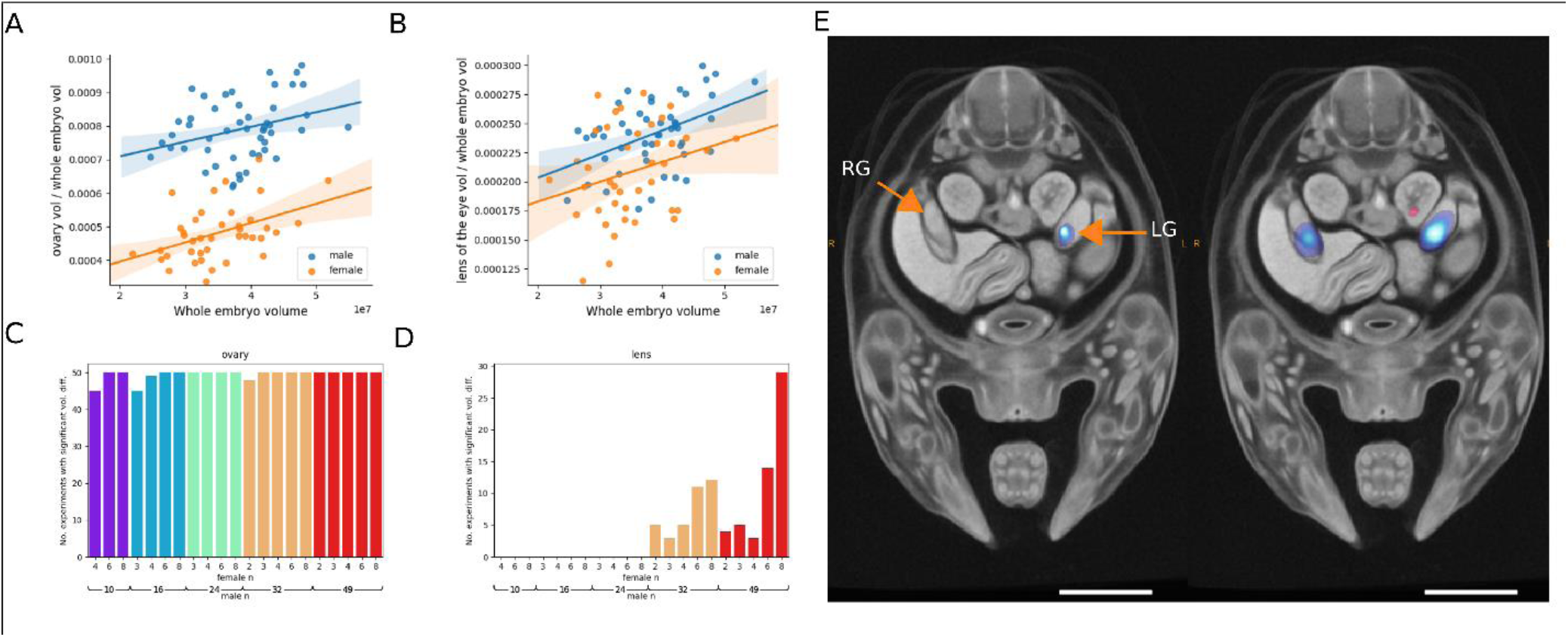
Identification of sex differences in wild type E14.5 mice. (A) A series of statistical tests were carried out with various combinations of male and female wild type sample size (with females relabelled as ‘mutant’). Each test was repeated 50 times with randomly selected specimens. The values reported are the number of times the gonad volume was reported significantly different (p < organ p threshold) in each set of tests. (B) Example results of Jacobian determinant analysis using 8 females and 8 males (left) and 8 females and 46 males (right). Scale bar = 1mm. RG: right gonad, LG = left gonad.

### Automated identification of developmental phenotypes in E14.5 mice embryos

The initial aim of LAMA is to automatically identify dysmorphology from data generated by IMPC and other projects To demonstrate its effectiveness on IMPC-generated data, we have chosen two exemplar mutant lines that illustrate its use in embryos with multiple dysmorphologies across the body and embryos with very specific, localised abnormalities.

The first example is *Wfdc2,* which encodes a protease inhibitor protein that is expressed in several tissues during mouse development prior to E14.5 (Lizio et al., 2015) including intestines, lungs and pancreas. *WFDC2* plays a role in cancer development (Bingle et al., 2002; Li et al., 2013) and two papers have recently shown *Wfdc2* homozygous mutants display severe pulmonary phenotypes in mice including collapsed lungs at perinatal day 1.5 (P1.5) (Nakajima et al., 2019), alveolar abnormalities, dyspnea and reduced blood oxygen saturation at birth (Zhang et al., 2019), but are otherwise anatomically normal. The IMPC reported a partially penetrant preweaning lethality phenotype for this line with 5.5% of the genotyped pups determined as homozygous for the mutation. Our analysis of four E14.5 *Wfdc2^-/-^* specimens resulted in significantly smaller organ volumes for the bronchi at the gene-level. (Fig 5A) but no significant specimen-level differences were observed for this gene. The Jacobian determinant analysis identified two significant regions that largely overlap with one of the bronchi after the FDR-corrected p-value threshold was raised to 0.1. This means that for this gene, the whole organ statistics were more sensitive than the Jacobian determinants. In all four mutants, the trachea and bronchus are visibly smaller in diameter (Fig. 5C-D), but otherwise appear normal.

**Figure 5.**
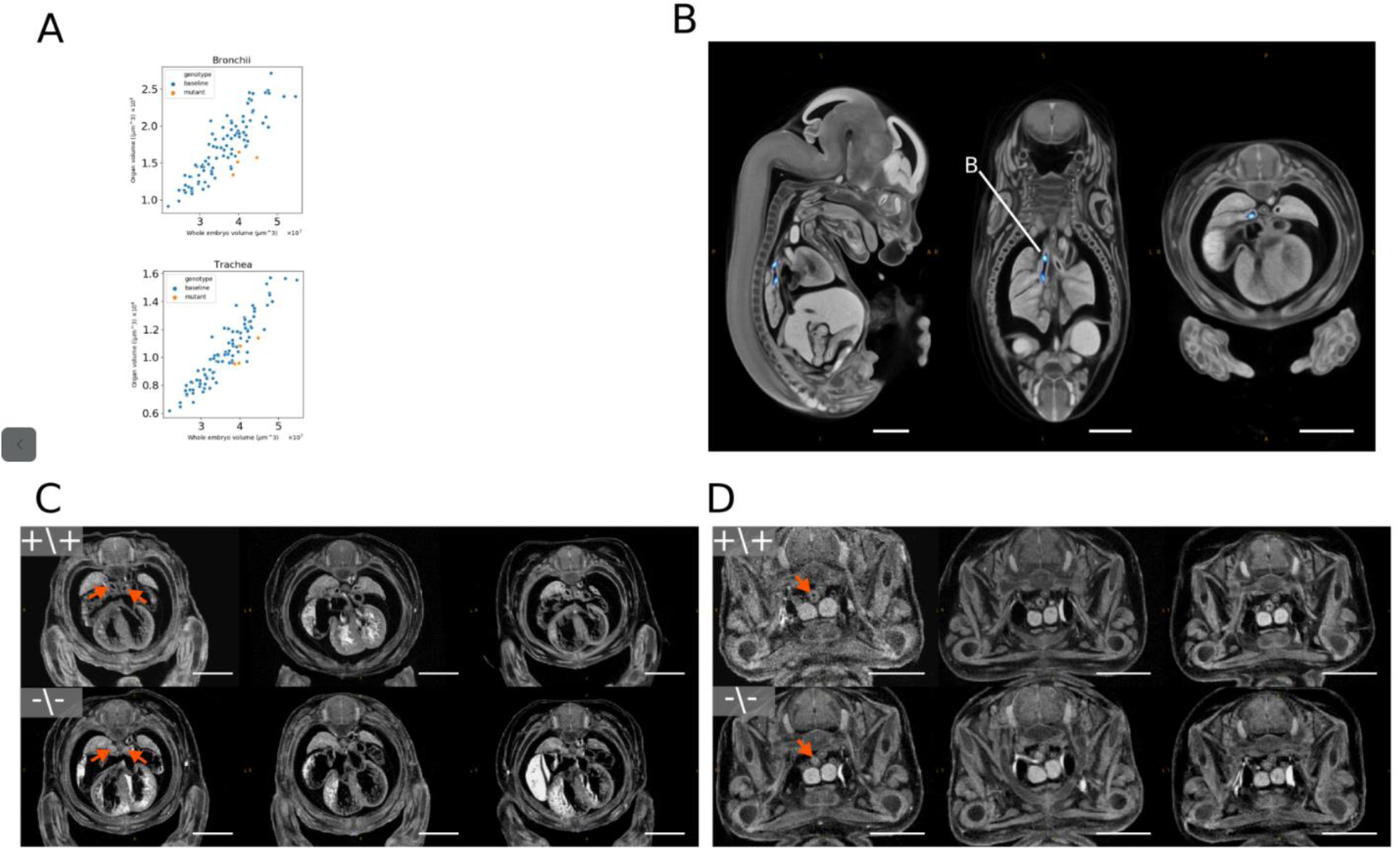
Automated identification of pulmonary system phenotypes in a *Wfdc2* knockout mouse line. (A) Organ volume plots from statistically significant organs. (B) Jacobian determinant analysis t-statistics (FDR corrected to q < 0.1) overlaid on the E14.5 population average. Blue regions indicate smaller bronchi in the mutants. (C-D) Axial slices of rigidly-aligned *Wfdc2* mutants (bottom) and whole embryo volume-matched wild type rigidly-aligned specimens (top). Arrows indicate the location of affected organs (C: bronchi, D: trachea). Scale bar = 1mm.

**Figure 6.**
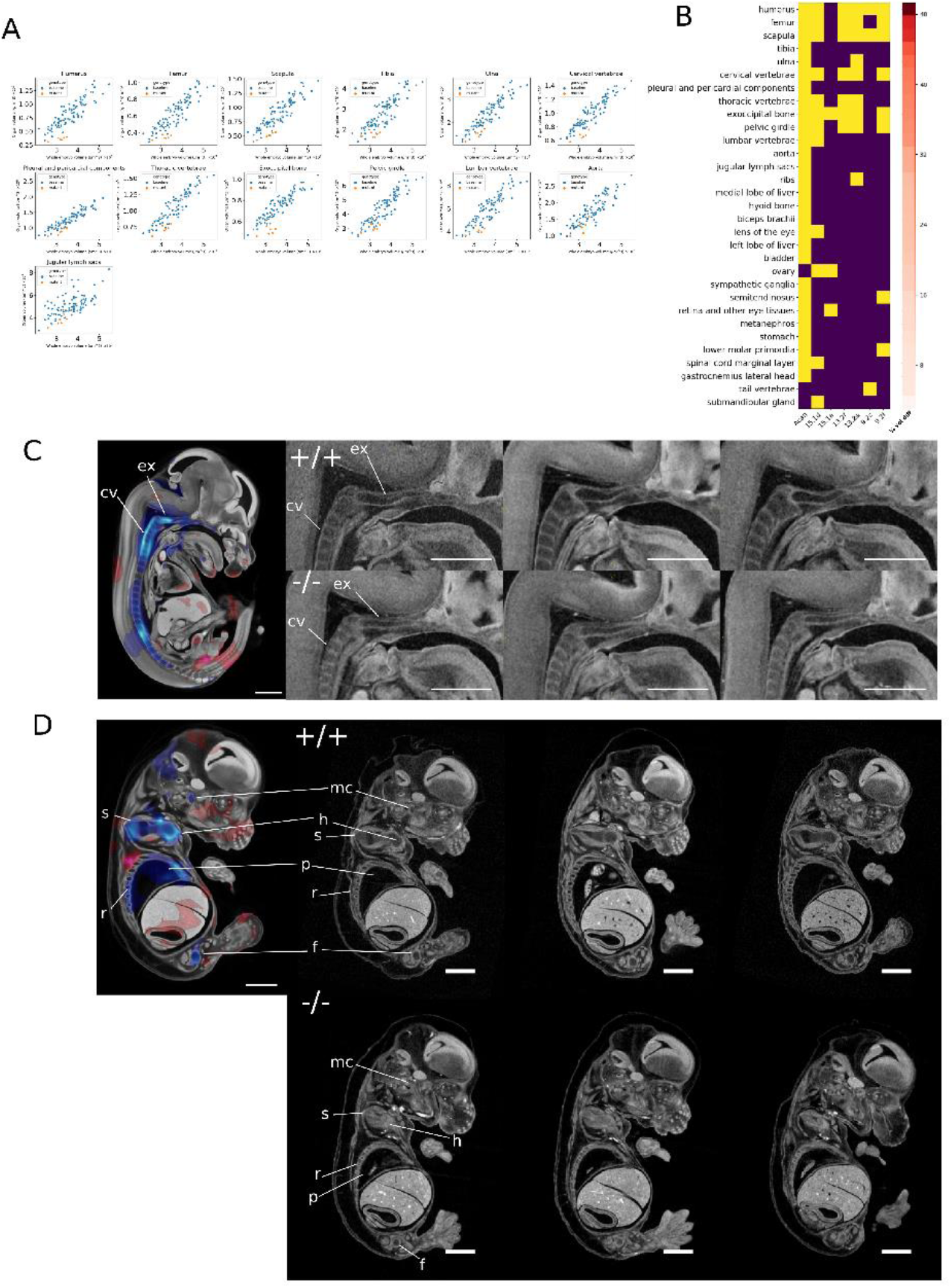
Analysis of *Acan*^*-/-*^ mutants by LAMA. (A) Scatter plots showing organ volume in voxels vs whole embryo volume in voxels for organs with a significant genotype effect. Only organs with a mean whole embryo-normalised volume difference, between controls and mutants, of > 15% shown (B) Comparison of line-level (*Acan*) and individual specimen-level results. Yellow cells indicate statistically significant organ volume differences for the genotype effect. The column ‘% vol diff’ indicates the mean organ volume difference (whole embryo normalised) between wild type and *Acan*^*-/-*^ mutants (C-D) Illustrative 2D sagittal slices highlighting identified dysmorphology. Left panels: Jacobian determinant t-statistics (FDR corrected to q < 0.05) overlaid onto the E14.5 population average. Right panels: rigidly-aligned *Acan*^*-/-*^ specimens (bottom), and stage-matched wild type rigidly aligned specimens (top) Scale bar = 1mm Legend: mc: Meckels’s cartilage, ex: exoccipital bone, cv: cervical vertebrae, h: humerus, s: scapula, p: pleural and pericardial components, r: ribs, f: femur

*Acan* encodes for the protein aggrecan, which is the primary proteoglycan in articular cartilage, is present in the extracellular matrix of long bone epiphyseal growth plates, and is required for normal bone development. *Acan*^-/-^ mice exhibit phenotypes associated mainly with abnormal bone morphology, including long bones, abnormal ribs and vertebrae as well as enlarged liver, and pulmonary hypoplasia (Table 3). Human diseases are associated with *ACAN* mutations including osteochondritis (Stattin et al., 2010) and skeletal dysplasia (Tompson et al., 2009). Along with complete preweaning lethality, the IMPC reports a reduced bone area composition and increased circulating cholesterol levels in adult heterozygous animals. We analysed six E14.5 *Acan*^*-/-*^ specimens with LAMA, identifying 28 statistically significantly organ volume differences. Of these, 13 had a mean volume difference between wild type and mutant of over 15% and are visible by eye (Fig. 6A), and 15 have a mean volume difference of below 15%, which makes visual identification of these differences difficult (Fig 6 suppl. 2). 12 of the significant organs are bones including all those present in the Mouse Genome Informatics (MGI) annotations (Table 3) (see Fig 6C-D for examples of identified dysmorphologies). Of these gene-level annotated organs, the specimen-level analysis assigned annotations to 15 of them (Fig. 6B) showing a considerable overlap between these two analyses. There were three significant organs highlighted by the specimen-level analysis that were not present at the gene-level, representing just four individual calls out of a total of 42 specimen-level calls, including one tail vertebra annotation. The significant Jacobian determinant voxels indicate a smaller Meckel’s cartilage in the mutants, which is not identified in previous literature or by our organ volume difference test, but is visibly different in the mutants (Fig. 6D). We did not find a significantly smaller lung volume difference for any of the lung lobes, which would indicate hypoplasia as previously reported (Houghton et al., 1989), but the inverted lung lobe labels indicate an acceptable registration accuracy (Fig 6 suppl. 1). The position of the lungs within the thoracic cavity however looks altered, possibly due to changes in the thoracic cavity size. We did not identify the remaining previously reported phenotype of tracheal cartilage morphology either.

**Table 3.**
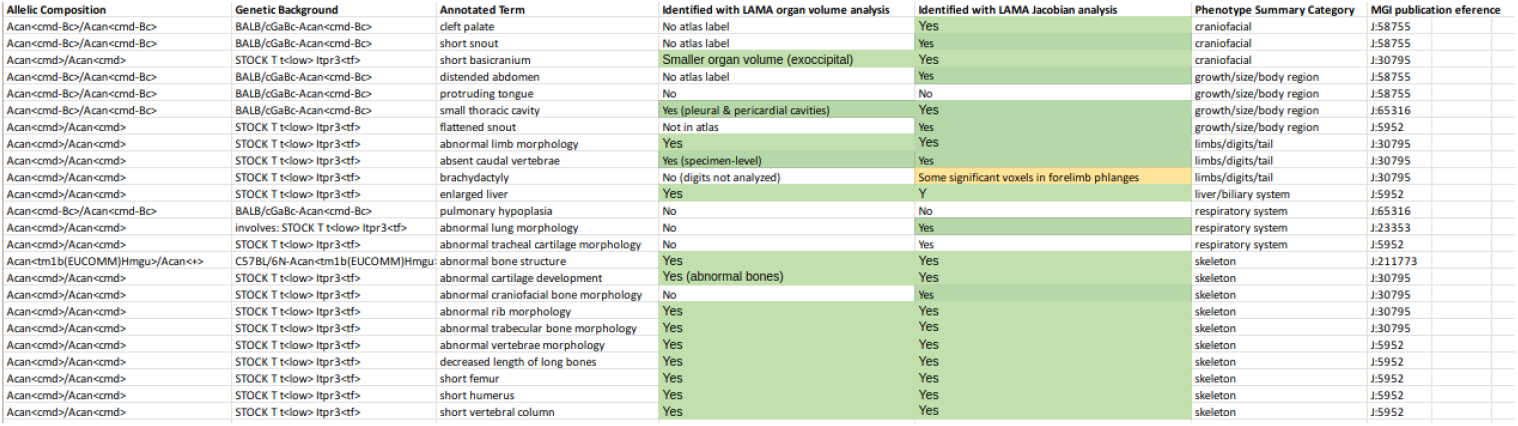
Phenotypes recorded for Acan ^-/-^ mutant mice in the MGI database. Green cells indicates concordance with recorded phenoty

## Discussion

In this work we present a new automated computational phenotyping pipeline for dysmorphology detection in mutant mouse embryos along with a novel, highly-detailed E14.5 anatomical atlas. The LAMA pipeline is easy to install and run, and uses fewer resources than currently available software. We have undertaken validation of the pipeline and provide data on the effect of developmental substage and sample size on phenotyping results, as well as showing that LAMA can uncover previously reported and new phenotypes from E14.5 IMPC knockout mice embryos. This pipeline can accelerate the automatic analysis of 3D embryo data at this developmental stage (and will be adapted for other stages) within large scale projects such as the IMPC and smaller challenge-led projects driving forward the use of disease models in scientific discovery.

We employ an image registration pipeline to spatially normalise wild type and mutants in a similar manner to a previously reported pipeline (Wong et al., 2012; Wong et al., 2014), but with the following two key differences. Firstly, during spatial normalization of the wild type and mutant images, LAMA uses the population average as the fixed image at each stage. This contrasts with Wong et al., (2012) where the wild type and mutant images (8 + 8) are spatially normalised by a combination of initial pairwise registrations in the linear stages and then non-linear registrations. In addition, the fixed image is the average intensity image from the previous stage. Thus, the specimens from each mutant line, and its controls, exist in their own unique coordinate space. By contrast, our approach allows the initial spatial normalization of all the wild type images into the population average space once only (as opposed to registering wild types for each mutant line), and to reuse the resulting data as controls for each mutant line, enabling us to dramatically increase the number of wild type embryos used as baseline comparisons. Secondly, we have shown that even after normalizing volumetric data by whole embryo volume, E14.5 developmental substage-specific organ volume differences are widespread across the embryo. These stage-dependent growth rate differences can lead to false positive results if the statistical model does include these substage data (Fig 3). Wong et. al., (2012) account for developmental substage variability by normalizing organ volumes and Jacobian determinants to whole embryo volume and report an organ volume standard deviation of 8-13% amongst wild type E15.5 embryos. In the current study, we sought to include as many wild type controls as possible in order to increase the power of our analyses, which increases the range of developmental substages in our control set, and we have adopted a modified linear regression model that controls for this confounding effect. Our choice of whole embryo volume (WEV), as a staging metric was motivated by the ease by which it can be calculated after embryo spatial normalization, and should correspond well to crown-rump length and whole body weight, which have been used previously to stage embryos (Dagg 1963; Peterka et al., 2002). Alternative methods that rely on the appearance of external features of the embryo (Theiler et al., 1989; Boehm et al., 2011; Geyer et al., 2017), may be more accurate, but these are not immediately amenable to automated analysis at E14.5. A promising approach to automated staging involves spatially, and temporally normalizing embryos to a four dimensional population average model embryo, with developmental stage as the fourth dimension (Wong et al., 2015). However the temporal dimension of the reported 4D model, extends only to E14.0 and it is generated from optical projection tomography images, a different modality to that used in this project. During this analysis we identified 81 organs that show a statistically significant staging effect (Table 2) which likely reflects different rates of growth of various organs at different E14.5 developmental substages, and this represents the most detailed embryo-wide data of E14.5 substage-specific organ growth rates that we are currently aware of.

Males and females are sets of animals with convenient and specific gonad differences that enabled us to test the ability of LAMA to identify anatomical differences. We found that we were able to uncover statistically significant organ volume differences in the gonads while keeping false positives low. To assess the performance on mutant data, we tested LAMA on two IMPC-generated knockout lines. The first (*Wfdc2^-/-^*) was predicted to display specific pulmonary abnormalities and the second (*Acan*^*-/-*^), a mutation shown previously to display severely-dysmorphic phenotypes across the whole of the embryo. Our automated analysis of E14.5 *Wfdc2^-/-^* embryos revealed two significantly different organ volumes: those of the trachea and bronchi, which are novel findings for this gene. These overlap broadly with the locations of the previously reported pulmonary-specific abnormalities in *Wfdc2*^-/-^ mice, including the absence of mature club cells from the bronchi and trachea, postnatally-collapsed lung, reduced lung surfactant levels (Nakajima et al., 2019), and alveoli abnormalities (Zhang et al. 2020). The novel phenotypes bring forward the time when gross abnormalities due to loss of *Wfdc2* first become visible during embryo development (previously postnatally), and therefore add new temporal information to the role of *Wfdc2* in pulmonary development.

In addition, we have shown that LAMA can recapitulate the majority of the previously reported *Acan*^*-/-*^ phenotypes, which will greatly speed up the annotation of this severely-affected mutant. There were also four significant organ volume differences for *Acan*^*-/-*^ that have not previously been reported. It is possible that these are novel phenotypes, but it is difficult to confirm this by looking at the CT images. It is possible that the apparent abnormality is due to proximity to actual severe dysmorphology and during the registration process were warped along with the abnormal organ. For example, two of these organs without previous reports, sympathetic ganglia and the spinal cord marginal layer, are located close to the affected vertebra.

As efforts are underway to reduce the numbers of animals used in scientific experiments we wanted to test whether LAMA could identify dysmorphology with low mutant sample number. In addition, being able to use low sample numbers would let us investigate the effects of incomplete penetrance and variable expressivity as well as providing phenotype data from mutant lines where many specimens do not reach the developmental stage being tested. To begin to answer this, in the sex difference test we show that increasing the control sample size from 8 to 46 greatly increases the power to detect anatomical differences and that by using many controls it is possible to sometimes uncover phenotype information even with a low mutant sample of one. We also show that when applying the specimen-level organ volume analysis to the *Wfdc2^-/-^* and *Acan*^*-/-*^ mutant lines, we were able to generate annotations on individual specimens. These specimen level annotations mostly overlapped with those assigned at the gene-level providing support for these phenotypes. These results lead us to recommend the following for our high throughput automated screen of mutant mice: to increase sample size to include as many control specimens as possible, and that specimen-level analysis is feasible at least on a subset of organs, but likely those which are most severely affected.

The choice of registration parameters involves a compromise of balancing good registration accuracy on some organs with misregistration at others. We found that the gonad registration, for example, could be improved by removing much of the registration constraint, but this led to over-warping at the heart. One solution to this could be to use multiple sets of registration parameters, each optimised to different parts of the atlas. Another approach could involve methods that do not require co-registration of images that could be applied to organs with a high registration error rate. This approach has been previously addressed (Yan et al., 2017; Ashish & Brusniak 2018) but only on a limited number of organs, and these methods are yet to be applied to embryonic mice. LAMA is able to perform statistical analysis on the voxel intensities of the spatially normalised images, but we found that the image data used in this study contained large differences in intensity profiles that were possibly due to the different users and imaging equipment involved in image acquisition over a number of years. For this reason, we have concentrated our current analysis of mutant lines on organ volume differences and Jacobian determinant analysis, which are both more robust to varying intensity profiles. Future work may look towards employing more sophisticated image normalization methods and exploring the analysis of other image features, such as textures, that may be less susceptible to intensity profile differences. Future work will include the adaption of the pipeline to other key developmental stages. E18.5 is a key developmental stage that we are currently working towards as it is important for the analysis of gene mutations that result in perinatal lethality and subviability. Earlier stages may also be amenable such as E12.5, but the earlier stages such as E9.5 may prove difficult for a registration-based approach due to the rapid developmental changes at this time point and the fact that mutations that cause lethality at this stage cause very extreme dysmorphology.

To conclude, we provide a new, easy-to-use pipeline for the automated analysis of E14.5 mouse embryos, along with a highly-detailed atlas that will be useful for both manual and automated studies of mutant mice. We have provided information on the differential growth rate of organs within the E14.5 developmental stage, shed light on the optimal number on the samples to use in mouse embryo phenotyping studies and have shown that LAMA can detect sex-specific differences and abnormal anatomy in E14.5 mutant embryos.

## Materials and methods

### Mice

All animals were housed and maintained in the Mary Lyon Centre, MRC Harwell Institute under specific pathogen-free (SPF) conditions, in individually ventilated cages adhering to environmental conditions as outlined in the Home Office Code of Practice. The Acan^tm1b^ allele was obtained by cre deletion of C57BL/6N-Acan^tm1a(EUCOMM)Hmgu^/H

(EM:10224) mice as described in (Birling et al., 2019). Homozygous mutants are named *Acan*^*-/-*^ here. The C57BL/6NTac-Wfdc2^em1(IMPC)H^/H (EM:11407, homozygous mutants named *Wfdc2^-/-^* here) was obtained by genome editing as described in (Mianné et al., 2017). Lines were maintained by crossing heterozygous animals with inbred C57BL/6N wildtype animals. Mice were euthanised by Home Office Schedule 1 methods.

### Micro-CT imaging of whole embryos

14.5 days post coitum (E14.5) female mice were sacrificed by cervical dislocation and the uterine horns removed into ice-cold phosphate buffered saline (PBS). Embryos were extracted and a piece of yolk sac collected for genotype analysis. Embryos were fixed in 4% paraformaldehyde (PFA) at 4°C and left overnight. After fixation, the samples were washed and stored in PBS at 4°C. For staining, samples were rinsed in dH_2_O for 10 minutes before being submerged in 50% Lugol’s solution and protected from light. Embryos were then left in the contrast agent for 2 days. Following staining, embryos were washed in dH_2_O for at least one hour, embedded in 1% agarose (in dH_2_O) and left at room temperature for a minimum of two hours.

High resolution micro-CT images (SkyScan 1172, Bruker) of agarose-embedded embryos were acquired at a source voltage of 70 kV, with the current set at maximum (∼100 mA). Specimens were imaged, in a standard orientation, at 3 μm with a 0.5 mm aluminum filter. X-ray projections were acquired at 0.25° increments, and reconstructed using the Feldkamp algorithm (Feldkamp et al. 1984) provided by NRecon (Bruker). Ring artefact corrections were applied as necessary. Reconstructions were automatically cropped to remove background and scaled to 14 um isotropic voxels using the HARP software (Brown et al., 2016).

### Registration pipeline implementation

The image registration pipeline was written in the Python programming language (Python 3.6), adapting a modular design that allows for individual components (registration, inversion, statistics etc.) to be run either sequentially or independently using simple TOML configuration files. Individual image registrations are performed using the elastix toolkit (S. Klein *et al.*, 2010; Shamonin *et al.*, 2013). The linear model analysis is implemented in R. All code is available on Github (https://github.com/mpi2/LAMA/releases/latest) and is tested to work on Ubuntu 18.04, and Windows. To make the installation of LAMA as easy as possible and to help data reproducibility, LAMA is available via the PyPi Python package repository (lama-phenotype-detection).

### Population average construction

Micro-CT images from 14 specimens of mixed sex were used in the creation of the population average through a multi-stage and resolution registration process. The initial fixed image was chosen at random from the input images and all other images are rigidly-registered towards it. The registered output images were averaged creating a rigidly-aligned blurry average. The outputs from the rigid registration are affinely-registered towards the blurry average to account for differences in overall size. The blurry affine average from this step was then used as the fixed image for deformable registration, which allows non-linear transformations. The deformable B-spline registration process contained five resolutions with decreasing control point spacing and blurring at each resolution, with a final grid spacing of 8 voxels, sequentially aligning smaller anatomical structures (Fig. 1A). The parameter file for our population average can be found here (https://github.com/mpi2/LAMA/configs/301015_pop_avg.toml) (link to population average when available).

### Image segmentation/E14.5 Atlas creation

Key anatomical structures within the E14.5 population average were identified manually by referencing the online digitised mouse atlas (Graham et al., 2015), and which itself is based on *The Atlas of Mouse Development* (Kaufman, 1992). Moreover, structures that could be identified were restricted to those that showed good contrast and resolution within the population average. ITKSnap was used (Yushkevich et al., 2006; www.itksnap.org) to create the segmentations, and dependent on size and complexity of an anatomical structure, were produced using semi-automated and manual methods and combined into a single label file until 184 structures were segmented at E14.5. These structures were then merged with some previous segmentations of brain structures derived from an E15.5 atlas (Wong *et al.*, 2012), to give a total of 184 anatomical components. Small, spindly labels in the atlas were flagged by taking the mean of a distance transform for each label, and flagging labels with a value < 1.5. The atlas and associated metadata file are available (link when available).

### Generation of data for phenotype detection

Baseline and mutant specimens are registered onto the previously created population average. The outputs of this registration include the non-affine spatial transformations, co-registered images and the Jacobian determinants det(J_F_), a scalar field which describes the local volume change in each voxel. Statistical analysis of these outputs (described below) produce statistical parametric heat maps that can be overlaid onto the population average applying the computed spatial transformations, or superimposed onto the input images applying the inverse of the spatial transformations. The statistical parametric heat maps can be viewed with the Volume Phenotype Viewer (VPV) described previously (Brown *et al.,* 2016; https://github.com/mpi2/vpv)

A typical project involves generating baseline data from wild type animals, before running all the mutant lines. For each line, the mutant and all the baseline data at each voxel, within a mask region, are fitted to a linear model with genotype and staging metric (whole embryo volume) as fixed effects, returning a t-statistic and p-value for the genotype effect.

## Statistical analysis

Multiple linear regression analysis was conducted in R (R Core Team, 2018) using the lm function from the MASS package (Venables & Ripley, 2002). Benjamin-Hochburg FDR (BH-FDR) correction was done using the padjust R package (Benjamin & Hochburg 1995).

### Voxel-level data

Jacobian determinants, generated at each voxel within the population average mask, provide information about how the registration has behaved at discrete locations in an image. The scalar value of the Jacobian determinant at a given location is the factor by which that region has expanded or shrunk in volume during registration. This approach, known as tensor-based *morphometry*, can be used to reveal biologically significant localised shape or size changes within a population (Ashburner & Friston, 2000). As well as the Jacobian determinants, the raw registered intensity images are analysed. To account for image intensity differences between the input images, LAMA includes an optional image intensity normalization step. Images are linearly normalised to the mean intensity of voxels across all the inputs or alternatively a region of interest (ROI) specified in the target coordinate space. To account for small registration inaccuracies, a Gaussian blur of full-width-half-maximum (FWHM) 100 μm is applied to voxel-level data. Each voxel is fitted to a linear model (*R* notation: *voxel ∼ genotype + whole embryo volume*). We use whole embryo volume as a proxy for developmental stage, and the addition of it as a fixed effect models out changes that are due to developmental stage only. To account for multiple testing, the resulting p-value maps are corrected using the Benjamini Hochberg method. The final parametric heat maps are made by thresholding the t-static volume at q>0.05. This is output as a volume that can be overlaid onto the target or registered image in VPV.

### Whole organ volume analysis

The atlas is inverted back onto the rigidly-aligned volumes, after registration, which effectively segments these volumes. These segmentations are used to calculate organ volumes for each specimen as well as providing a visual atlas for identifying structures in the original inputs and determining registration accuracy. The organ volume data is much less complex than the voxel-based data (the number of organs vs number of voxels) allowing us to employ a more robust method for multiple testing correction described previously by Hrabě de Angelis et al., (2015). To summarize, organ-specific null distributions are generated by sampling synthetic mutants from the baseline data in such a way as to match the distribution of mutant specimens per line. Synthetic mutants are fitted to a linear model as described previously (*organ volume ∼ genotype + whole embryo volume*) and the genotype effect p-values are stored. Alternative distributions are made by collecting genotype effect p-values from testing the real mutants of each line. To obtain a data set-wide p-value threshold per organ, p-values for the organ are ranked and a descending p-value search is conducted starting at p=0.05 until a value is found where the proportion of alternative p-values under the threshold divided by the proportion of null p-values is < 0.05. Mutant p-values below this threshold are assigned as significant, which sets the organ-specific FDR to 5%.

### Developmental Substage analysis

To simulate the effect of analysing mutant lines containing specimens at early or late E14.5 developmental substage, two datasets were created from 99 wild type specimens by relabeling subsets based on the calculated whole embryo volume (WEV) of each specimen. In the first set, eight specimens with the lowest WEV were relabelled as mutant. In the second set, eight specimens with the highest WEV were relabelled as mutant. To test whether a substage effect could be detected, the organ volume or Jacobian determinants data points from each set were then analysed by using the linear regression models (*organ volume ∼ genotype*) or (*Jacobian determinant ∼ genotype*). To test whether including WEV as a fixed effect could remove the substage confound, the organ volume or Jacobian determinants from each set were then analysed by using the linear regression models (*organ volume ∼ genotype + WEV*) or (Jacobian determinant ∼ genotype + WEV). The resulting organ volume, or Jacobian determinant, genotype effect p-values were corrected for multiple testing using the Benjamini Hochberg method.

To assess the contribution of developmental substage on local relative volume differences across the embryo, Jacobian determinants or organ volumes from 99 wild type specimens were analysed by using the linear regression model *(organ volume ∼ whole embryo volume*). The resulting organ volume, or Jacobian determinant, genotype effect p-values were corrected for multiple testing using the Benjamini Hochberg method.

### Detection of sex-specific differences

The spatially normalised data from 93 wild type specimens were split by sex. For each experiment, a different number of males and females were chosen at random without replacement and the females were relabelled as mutants. These data were then analysed by using the linear model (*organ volume ∼ sex+ whole embryo volume*). For organ volume analysis, organ-specific p-value thresholds were generated using the permutation based method as described above. Only experiments where combinations of male and female sample size allowed at least 500 unique null permutations were included. Each experiment was repeated 50 times to gain a robust measure of the effect of sample number. For each set of experiments, the organs that were reported as having a significant organ volume difference were recorded.

For the Jacobian determinant analysis q-values were generated from the resulting p-values using the Benjamini Hochberg method and threshold to q < 0.05. False positive rates for organs other than gonad or lens of the eye was calculated by dividing the total number of times a given organ passed the organ-specific significance threshold by the total number of tests performed.

### Variable penetrance and low *N*

To identify potentially variable expressivity or incomplete penetrance of organ volume phenotypes, specimens were treated as for the line-level analysis above. With the number of mutants reduced to one the FDR threshold was increased to 20%. The voxel-level data is similarly processed as the line-level voxel data and the voxels are thresholded to an FDR of 5%.

## Optimization and quality control

At each stage of the registration process, the image similarity metric output by elastix is plotted against iteration number for each spatial resolution allowing the user to select an optimal number of iterations per stage.

Image registration can sometimes fail to produce acceptable results, for example when the moving image is over fitted to the fixed image, producing unrealistic warping of images. To check for issues such as these, after each registration stage an additional HTML report is generated which contains mid-sagittal slices for each registered image for rapid quality control. Another issue that can be encountered using bspline-based image registration is folding of the deformation field which prevents topology preservation, reducing the accuracy of the propagation of labels from the atlas to the specimen images. This problem can be detected by the presence of negative Jacobian determinants. In the case of negative Jacobian determinants, instead of outputting log-transformed Jacobian determinants, only the negative Jacobian determinant regions are displayed, allowing the user to quickly identify problematic regions within the registered images. The registration stage of the analysis is the most time-consuming part of the pipeline, and so it is a requirement to be able to optimize registration parameters, especially within a high throughput context.

### Comparing phenotypes to known phenotypes

Tables of known phenotypes were generated by querying MGI phenotype pages for the gene of interest (*Acan*: www.informatics.jax.org/marker/phenotypes/MGI:99602, *Wfdc2*: www.informatics.jax.org/marker/phenotypes/MGI:1914951). Only phenotypes generated form a homozygous null strains were kept to aid in the comparison. Duplicate and redundant phenotypes (e.g. *abnormal bone structure* if more specific bone phenotypes were present). Also removed were phenotypes that might not translate to a gross anatomical dysmorphology that could be potentially detected by LAMA (e.g. deafness).

## Figures

**Figure 4, supplementary 1.**
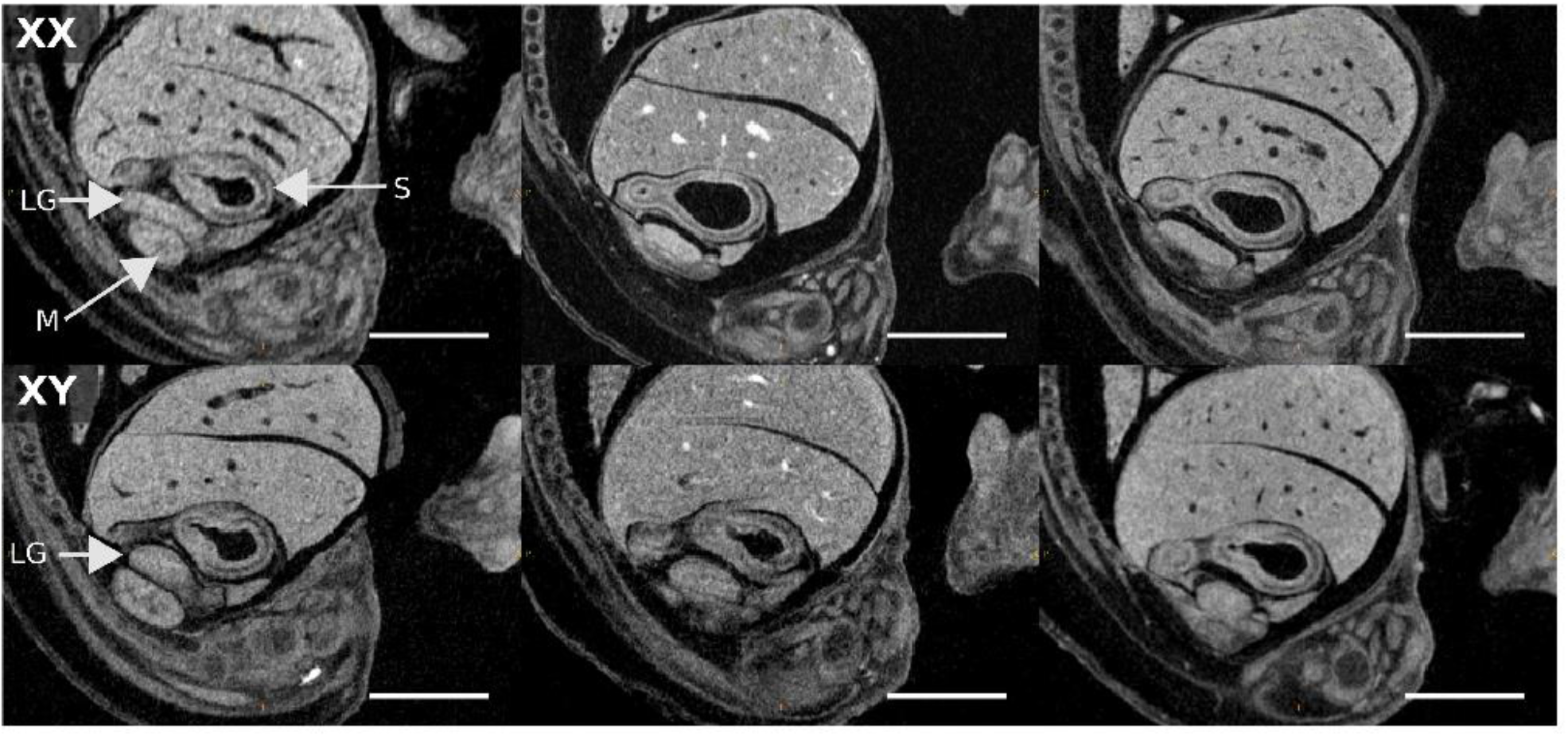
Wild type embryos gonad examples. Sagittal sections of rigidly-aligned wild type embryo images showing the left gonad (LG). The stomach (S) and metanephros (M) are also indicated. Females (top) and males (bottom). Scale bar = 1mm.

**Figure 4, supplementary 2.**
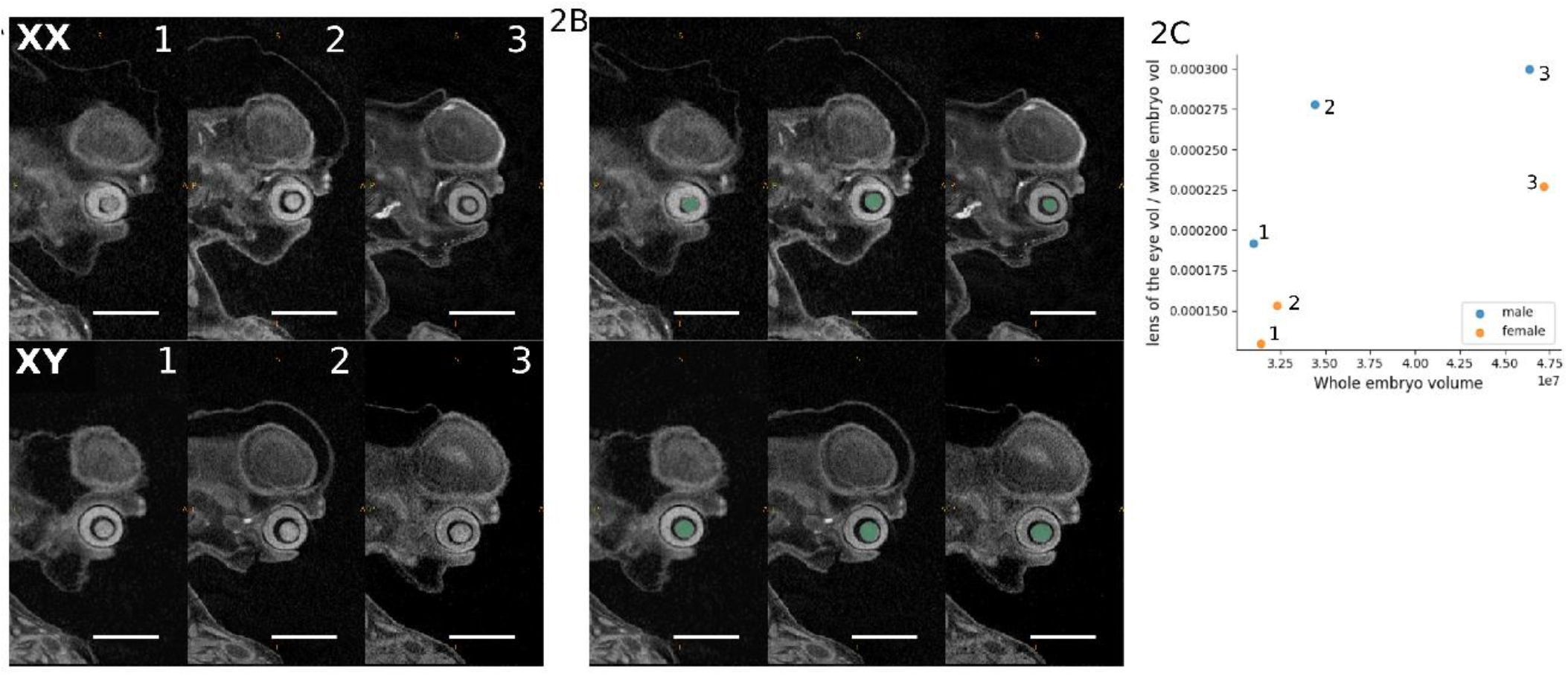
Differences in wild type lens volume. Representative sagittal sections of affinely-registered (normalised for overall embryo size) wild type specimens illustrating difference in lens sizes between females (top rows) and males (bottom rows). (A) Unlabeled affinely-registered images (B) Affinely-registered images overlaid with inverted lens label (green). (C) Plot of whole-embryo normalised organ volume against whole embryo volume.

**Fig 6. Supplementary 1.**
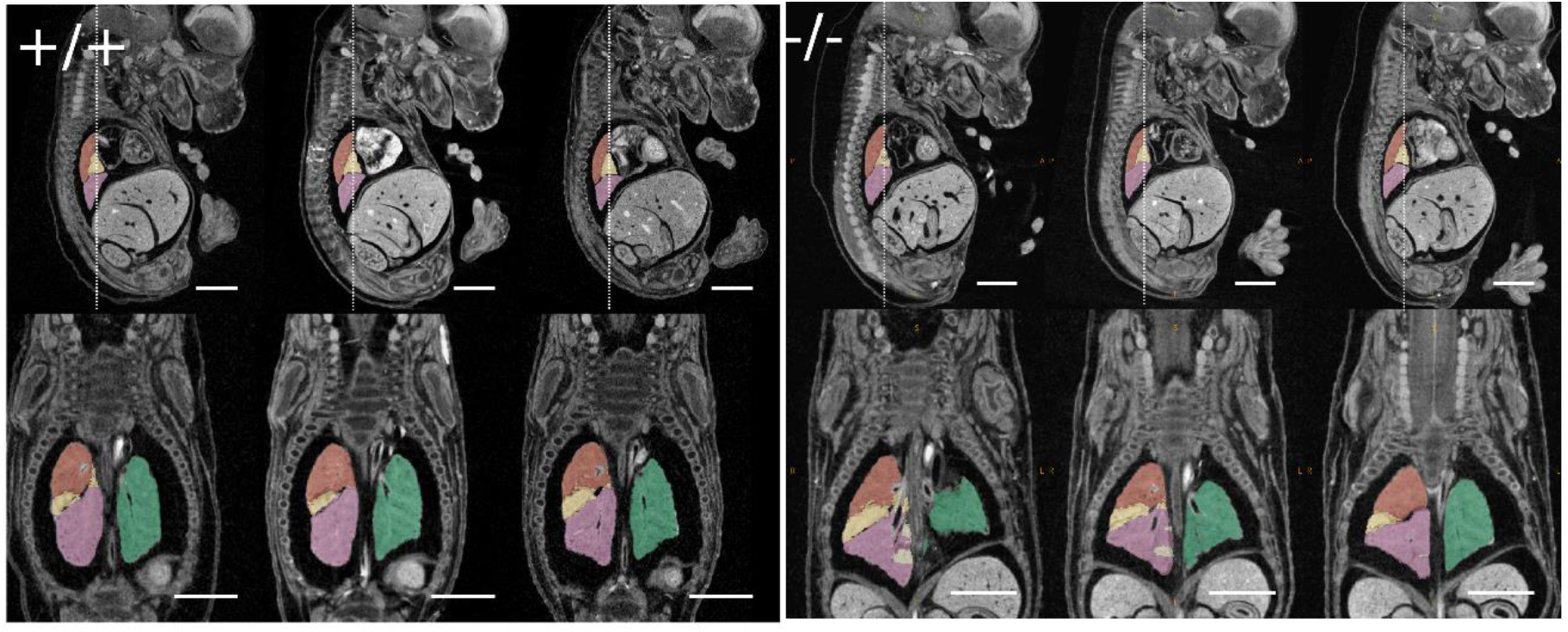
Comparison of wild type and Acan-/-mutant lungs. Rigidly-aligned specimens overlaid with individual lung lobe labels inverted from the atlas showing mutants (left) and stage-matched wild types (right). Top row are sagittal sections, bottom row are coronal sections. The dotted line in the sagittal sections corresponds to the section chosen for the coronal view.

**Fig 6. Supplementary 2.**
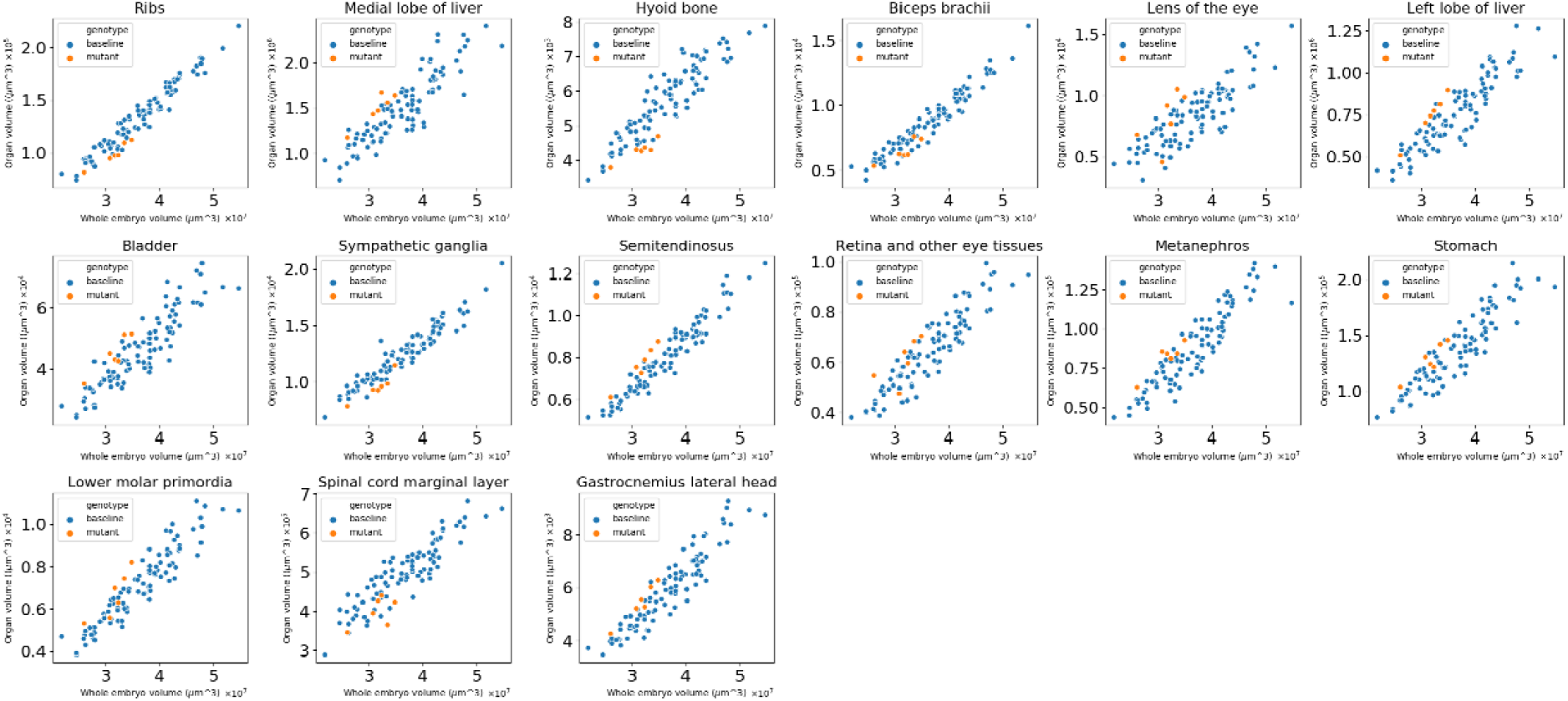
Significant organ volume differences in *Acan*^*-/-*^ null mutants. Scatter plots showing organ volume in voxels vs whole embryo volume in voxels for organ with a significant genotype effect. Organs highlighted here have a mean whole embryo-normalised difference of less than 15% between wild types and mutants. Blue markers are wild type controls, orange markers are mutants.

## Acknowledgements

We would like to thank James Cleak and and Zsombor Szoke-Kovacs for generating image data and helpful discussions. Also thanks to the husbandry team from the Mary Lyon Centre for generation and maintenance of all the mouse specimens used in the study.

